# Laboratory *An. gambiae s.l*. mosquito colonies show sustained high transmission of Microsporidia sp. MB and a small fecundity cost

**DOI:** 10.64898/2026.02.24.707712

**Authors:** Beth Crawford Poulton, Deepak-Kumar Purusothaman, Abdelhakeem Ismail Adam, Saré Issiaka, Ewan R. S. Parry, Roland Pevsner, Thomas H. Ant, Etienne Bilgo, Abdoulaye Diabaté, Steven Paul Sinkins

## Abstract

Microsporidia sp. MB, a microsporidian symbiont found naturally in *Anopheles* mosquitoes, has potential as a novel malaria control tool since it can inhibit *Plasmodium* development and transmission. The most feasible MB-based *Plasmodium* control strategy would involve dissemination through live mosquito releases, or release of spores infective to mosquito larvae. To implement either strategy, establishment of stable mosquito colonies carrying MB at a high frequency is likely to be essential. The progeny of field caught *An. gambiae s.l* from Burkina Faso were isolated for individual egg laying and tested for MB. The progeny of the MB positive females were pooled and this process was repeated for multiple generations. The relative density of MB in different life stages and tissues of the *An. coluzzii* host was examined using a novel duplex qPCR assay. We also examined the impact of MB on fecundity through individualization for egg laying and counting of eggs. Finally, we examined laid eggs for presence of MB spores. Three *An. coluzzii* colonies and one *An. gambiae s.l* hybrid colony were established with high prevalence and density of MB and were maintained for more than two years with minimal intervention. MB prevalence and density was highest in eggs and adult females and lowest in L4 larvae; in adults density was highest in the gonads. Additionally, MB density increased in ovary samples following blood feeding which was likely due to the activation of sporogony. The production of spores is the reason why MB-carrying females lay more white non-hatching eggs and show a small reduction in fecundity. Establishment of several stable MB carrying *An. gambiae s.l* colonies and understanding the impact of spores on fecundity are significant steps forward in developing MB as a malaria control tool.

## 1 Introduction

There were an estimated 263 million malaria cases and 597,000 deaths in 2023 alone (WHO, 2024) and despite progress in its control malaria remains a very serious global health challenge. The use of insecticide treated bed nets and indoor residual spraying of insecticides substantially reduced the number of malaria cases and resulting deaths between 2000 and 2015 (Bhatt et al., 2015), however this progress stalled with both case numbers and deaths being greater in 2024 than in 2015 (Venkatesan, 2024). The reasons for this are multifaceted but are largely due to the emergence of insecticide resistant mosquito populations (Ranson and Lissenden, 2016, Bilgo, 2024). It has become apparent that new control measures will be necessary if the burden of malaria is to be further reduced (Bilgo, 2024, Venkatesan, 2024).

Certain mosquito endosymbionts have been shown to interfere with pathogen development and transmission (Pumpuni et al., 1993, Bian et al., 2010, Bongio and Lampe, 2015, Ant et al., 2018). Therefore, they can be used as novel agents to control mosquito-borne diseases (Iturbe-Ormaetxe et al., 2011, Ricci et al., 2012). One approach that has proven successful for controlling dengue transmission by *Aedes aegypti* mosquitoes is the introduction and spread of the *Wolbachia* endosymbiont (Hoffmann et al., 2014, Nazni et al., 2019). Currently, two *Wolbachia* strains (*w*AlbB and *w*Mel) that were introduced into *Ae. aegypti* are being deployed for dengue control, and communities in countries such as Australia, Malaysia, Brazil, and Colombia have conducted well-received large-scale *Wolbachia* releases (Hoffmann et al., 2014, Nazni et al., 2019, O’Neill et al., 2019, Velez et al., 2023, Simmons et al., 2024). *Wolbachia* effectively prevents arbovirus transmission when introduced into *Ae. aegypti*, but no *Wolbachia* strains have been detected naturally in this mosquito species (Ross et al., 2020). The *Wolbachia*-carrying *Aedes* lines are derived from *Aedes albopictus* (*w*AlbB) and *Drosophila melanogaster* (*w*Mel) (Xi et al., 2005, Walker et al., 2011). Recent reports indicate the presence of naturally occurring *Wolbachia* in *Anopheles* mosquitoes (Baldini et al., 2014, Shaw et al., 2016, Niang et al., 2018). However, they occur at very low frequencies/intracellular densities, and their vertical transmission rates are relatively low (Gomes et al., 2017, Gomes and Barillas-Mury, 2018).

A naturally occurring microsporidian symbiont Microsporidia sp. MB (henceforth MB), was identified in *Anopheles arabiensis*, and was shown to inhibit *Plasmodium falciparum* development in the mosquito vector (Herren et al., 2020). MB was first detected in *An. arabiensis* and *An. Gambiae s*.*s* in Kenya (Herren et al., 2020, Nattoh et al., 2023) and has since been found in *An. coluzzii* in Niger (Moustapha et al., 2024) and in both *An. gambiae s*.*s* and *An. coluzzii* in Ghana (Akorli et al., 2021), Benin (Ahouandjinou et al., 2024, Tchigossou et al., 2025) and Burkina Faso (Millogo et al., 2024, Bilgo et al., 2025). MB has not been reported to be associated with significant morbidity or mortality in naturally infected mosquitoes. Transmission of MB appears to be impacted by several environmental factors often related to seasonality (Tchigossou et al., 2025) and water quality (Akorli et al., 2024), specifically temperature, humidity and diet (Boanyah et al., 2024, Bilgo et al., 2025, Otieno et al., 2025). Until recently, vertical transmission has been assumed to be the primary route for MB in mosquitoes (Nattoh et al., 2023, Onchuru et al., 2024, Makhulu et al., 2024, Parry et al., 2025) and (in *An. arabiensis*) sexual transmission from male to female a far less efficient secondary route (Nattoh et al., 2021, Maina et al., 2025). However, identification of octagonal sporogony in developing eggs post blood feeding (Parry et al., 2025, Herren, 2025) suggests a fork in the lifecycle which could play an important role in MB transmission although what these spores infect is not yet understood.

If MB is to be used for malaria control, it will be necessary to increase the proportion of mosquitoes carrying the symbiont in the field. This could be accomplished through release of mosquito-infecting MB spores or release of MB colonized mosquitoes. Each of these strategies will likely require large numbers of MB colonized mosquitoes, therefore establishment of laboratory reared colonies with high MB prevalence is essential.

## 2 Materials and Methods

### 2.1 MB colony establishment and maintenance

#### 2.1.1 Field Collection

Adult mosquitoes were collected with an aspirator from Vallee du Kou, VK7 (11°23’N, 4°24’W) and Soumousso (11°04’N, 4°03’W) between 7 and 9 am from human and animal dwellings using the residual fauna capture method. Blood fed females were morphologically identified using the Gillies and De Meillon identification keys (Gillies and Coetzee, 1987) and transported to the Institut de Recherche en Sciences de la Santé (IRSS). The females from VK7 (G0) were blood fed (live rabbit) and their eggs (G1) were collected in one batch which was transported to the Centre for Virus Research (CVR), University of Glasgow.

Females collected in Soumousso were blood fed (live rabbit) and separated into cups to lay eggs individually. For each female that laid eggs, DNA was extracted using the Cetyltrimethylammonium Bromide (CTAB) method (Lardeux et al., 2008) and tested for MB by endpoint-PCR using established primer sequences (Herren et al., 2020) (Supplementary Table S1). Progeny from MB positive females were pooled, reared to adulthood as described below, blood fed (live rabbit) and eggs (G2) from these mosquitoes (G1) were collected as one batch and transported to the CVR in Glasgow.

For transport from IRSS to CVR, eggs were collected on damp filter paper which was then folded, enclosed in a Petri dish with damp cotton wool, and sealed with Parafilm®.

#### 2.1.2 Standardized laboratory rearing

In the CVR, all *An. gambiae s*.*l* mosquitoes were reared in insectaries with climate controlled to 25-28 °C and 60-80 % relative humidity with 12 h/12 h light/dark cycles. Adults were kept in BugDorm-1 or BugDorm-4 insect rearing cages provided 5 % sucrose solution ad libitum and a Hemotek artificial blood-feeding system (Hemotek Ltd) was used with either defibrinated horse-blood (E&O laboratories, UK & Thermo Scientific) or human blood (Scottish National Blood Bank) to induce egg production. Eggs were collected on wet, grade-1 filter-paper (Whatman plc) and hatched in distilled water with a pinch of bovine liver powder (Now foods). Larvae were reared in distilled water, fed with liver powder until the second instar then with TabMin fish food pellets (Tetra) until pupation, changing water 2-3 times during the larval cycle as necessary to ensure healthy development.

#### 2.1.3 Successful colony establishment method

After failing to acquire viable eggs from sibling crosses after the first blood feed in Glasgow for both VK7 (G1) and Soumousso (G2), males from an established laboratory *An. gambiae s*.*l* hybrid colony, ‘G3’ (Kindly provided by Dr. Gareth Lycett, Liverpool School of Tropical Medicine) were introduced to each cage and, after a third blood feed, enough eggs were collected to proceed with selection of MB positive colonies.

To maximize the likelihood of success, multiple oviposition’s were collected for 7 generations, leading to the establishment of several related but distinct colonies with high MB prevalence (Fig. 1). Some egg batches were collected for all females in a cage as described above for standard rearing – noted ‘pooled’ in Fig. 1. Other egg batches were selected for MB – the results of which are indicated in Fig. 1. Adult females three days post blood feeding were individualized for oviposition, followed by CTAB DNA extraction of females that laid eggs and PCR screening for MB. Only the progeny of MB positive females were pooled for the subsequent colony generation. MB Amp-V2 end-point PCR was conducted to screen G1-6, with validation using the existing MB specific primers MB18SF and MB18SR from Herren et al, 2020 (Herren et al., 2020) for G1-3. From ∼G7, the MB 18S (MT160806.1) and *An. gambiae* Elongation factor Tu (AGAP005128) multiplex probe-based RT-qPCR described below was used as the primary assay for all subsequent screening and experiments with only one exception. For only the BQ+ G9 selection a SYBR (Fast SYBR Green Master Mix, Thermo Fisher Scientific Life Technologies: 4385612) qPCR was used with MB 18S primers SP79_MB_QPCR1_F and SP80_MB_QPCR1_R, and control primers SP77_S7_F and SP78_S7_R. At G12 the primary DNA extraction method was changed from CTAB to grinding in 50-100 µL Sodium Tris EDTA (STE) buffer solution (Merck, 85810) and boiling at 95 ℃ for 10-15 min.

**Fig. 1:**
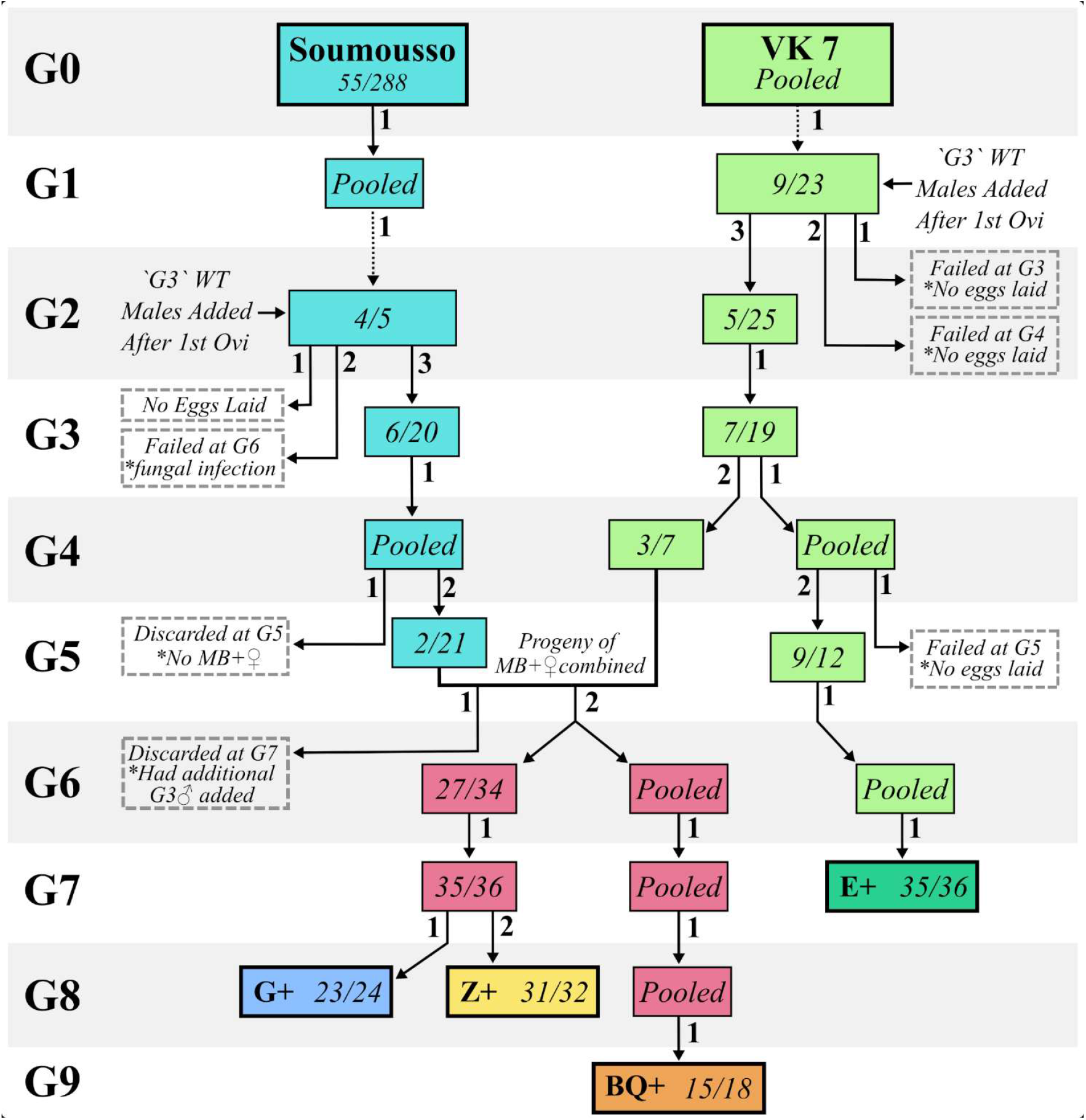
Initial Colony Establishment. Flow diagram indicating the process of establishing four Microsporidia sp. MB positive *An. gambiae s*.*l* colonies (G+, Z+, BQ+ and E+). Pooled indicates an adult cage for which eggs were collected in one batch where no selection occurred. X/XX indicates the number of females that were Microsporidia sp. MB positives / the total number of females that laid eggs and were screened molecularly at the generation (G) noted on the left. The small number beside each arrow indicates the oviposition number. Where an arrow splits this indicates an oviposition for which multiple cages of progeny were generated that were treated differently. The dotted arrows indicate the transfer of eggs from IRSS in Burkina Faso to CVR. Failed lineages are indicated with a dotted border.’G3’ WT refers to laboratory adapted ‘G3’ *An. gambiae s*.*l*. mosquitoes.

### 2.2 MB-18S plasmid construction

To validate new MB diagnostic PCRs, a plasmid carrying the MB 18S gene was produced. The MB 18S gene sequence was identified in NCBI (accession: MT160806.1), and amplified from our MB-infected mosquitoes, using the following primers SP35_Microspor-MB_Fwd and SP74_Microspor-Ubiq_rev-alt (Supplementary Table S1). The amplified PCR fragment was purified using agarose gel electrophoresis and cloned into the blunt vector backbone pJet1.2 using the CloneJet PCR cloning kit (Thermo Scientific) as per the manufacturer’s recommendations. The cloned plasmid was transformed using XL10-Gold® ultracompetent cells (Agilent Technologies). The purified plasmid was quantified using a Nanodrop spectrophotometer (Thermo Scientific) and plasmid copy numbers per µL were calculated using the Science Primer online tool (https://scienceprimer.com/copy-number-calculator-for-realtime-pcr). The plasmid map is provided in Supplementary Fig. S1 and the complete MB 18S plasmid sequence has been deposited in the NCBI database (accession: PX674407). The same primers were used to confirm the MB-18S sequence by sanger sequencing following colony establishment for each line (Supplementary Fig. S2).

### 2.3 MB Diagnostic methods

#### 2.3.1 New end-point PCR assay (Amp-V2)

A new end-point PCR was designed using the MB 18S scaffold sequences (shared by Dr. Lilian Mbaisi Ang’ang’o / Dr. Jeremy Herren, ICIPE, Kenya), targeting regions that were conserved within MB but exhibited some variation when compared to other closely related microsporidia (Supplementary Fig. S3). The primers SP118_MB_Ampv2_F and SP119_MB_Ampv2_R (Supplementary Table S1) were designed using the NCBI Primer-BLAST tool using the standard parameters. The sensitivity of the new MB 18S primers was tested using the MB plasmid serial dilutions from 1×10^0^ to 1×10^10^ copies and the Q5 High Fidelity Polymerase (New England Biolabs).

#### 2.3.2 MB duplex qPCR

To improve the specificity and efficiency of MB quantification, a multiplex probe-based real time qPCR assay was developed. For MB and 15 closely related species, the available sequences for the 18S ribosomal RNA gene were aligned using the MAFFT nucleotide multisequence alignment tool in Benchling (Supplementary Fig. S4). From this alignment, regions of low sequence homology between MB (MT160806.1) and other species were identified, and primers and probes were designed using the IDT Mismatch ToolTM and NCBI Primer BlastTM to maximize specificity to MB. Primers and probes were designed for the *An. gambiae* elongation factor (EF) thermo unstable (AGAP005128) gene (Supplementary Fig. S4). The final primer and Affinity Plus probe (including some locked nucleic acid bases) combination (Supplementary Table S1) was validated for PrimeTimeTM Gene Expression Master Mix (IDT) which was used as per manufacturer’s instructions with final concentrations of 0.2 µM and 0.5 µM of each probe (agEF_Probe and MB-18S_Probe) and primer (SP121_MB-18S_F, SP122_MB-18S_R, SP172_An.gam-EF_F and SP173_An.gam-EF_R) respectively with 1 µL template in a 10 µL total reaction volume. qPCRs were run in triplicate using a QuantStudio 3 or 5 on 96 or 384-well plates respectively.

Analysis of qPCR data was conducted in R (v4.1). Any samples where fewer than 2 technical replicates had AgEF_Probe Cqs that were > 32 or undetermined these samples were considered to have insufficient DNA quality and were removed from the analysis. MB positive samples (MB+) were defined as those for which have a MB-18S_Probe Cq < 35. Negative samples (MB-) were defined as those for which had Cq >35 or was undetermined for MB-18S_Probe. Pairwise proportion tests with Bonferroni correction were used to assess the difference in proportion MB+ between groups.

Relative density of MB_18S_Probe compared to AgEF (2-ΔCq) was calculated for each well individually and used to calculate the arithmetic mean density of technical replicates for each sample. Where mean density of groups of samples is presented (orange diamonds on plots) the geometric mean was calculated to account for the base 10 logarithmic transformation of the data which was used in statistical analyses. For each experiment, a one- or two-way ANOVA was conducted (Supplementary Table S2) followed by two-tailed pairwise estimated marginal means with Bonferroni correction for multiple comparisons using functions from the rstatix package (v0.7.2).

### 2.4 Short interspersed element-200 bp (SINE200) PCR

SINE200 PCR (Santolamazza et al., 2008) was used to determine the molecular forms of *An. gambiae* in different MB positive lines as described in the MR4 Methods in Anopheles Research manual (MR4 et al., 2015). 1 µL genomic DNA template and Q5 polymerase was used as per manufacturers instructions.

### 2.5 MB density determination

#### 2.5.1 Different life stages

To determine the MB density across the different developmental stages, a probe-based duplex qPCR assay was performed. Genomic DNA from the samples was extracted using the modified STE Buffer (Sigma Aldrich) protocol. The different life stages of mosquitoes assessed for MB density using MB duplex qPCR were eggs, larvae - first to fourth instar (L1-L4), male and female pupae, and male and female adults (3-7 days post emergence, virgin), with a sample size of n = 12 per life stage for each of three biological replicates. Sex of both adults and pupae were determined by visual examination of external genitalia at the pupal stage (Poulton et al., 2021).

#### 2.5.2 Dissected adult tissues

Salivary glands, gut, ovaries, spermatheca and the remainder of the carcass were dissected from “virgin unfed”, “mated unfed” and “mated blood-fed” adult female mosquitoes for STE buffer DNA extraction and MB duplex qPCR to test the MB density across both the somatic and germline tissues. Virgin females were sexed at pupal stage by visual examination of external genitalia (Poulton et al., 2021). Mated and virgin adult males were dissected, and the testis (including accessory glands), gut and the remaining carcass were separated for separate evaluation. The adult mosquitoes used for tissue dissections were three to seven days post eclosion except for blood-fed females which were dissected three-days after receiving a blood meal at three to seven days post-eclosion (before oviposition). DNA was extracted from individual dissected tissues using 50 µL STE Buffer and the relative density of MB was quantified using the MB duplex qPCR. For each group and tissue, 12 samples were collected for each of three biological replicates.

### 2.6 Fecundity Assessment

Mosquito egg-laying (fecundity) was assessed in the MB positive and negative Z lines. The Z-line was established through selection of the progeny of MB negative females which were identified during the selection of the Z+ colony. Three to seven-day-old pre-mated females were membrane-fed using human blood (Scottish National Blood Bank) on the Hemotek feeding system as per standard instructions. Only engorged females were selected on day three after a blood meal and individually placed in a 7 ml plastic vial (Greiner, 189170) for egg-laying. Each plastic vial was filled with 0.5-2 mL of distilled water and covered with a cotton ball. Three biological replicates were completed with positive and negative colonies assessed in parallel, and for each colony, 60-100 females were used. After 24 hours post-individualization, eggs were counted from each vial; both white and dark eggs were identified and noted separately. At the end of the experiment, females that laid eggs were collected individually for STE extraction and MB duplex qPCR to quantify the relative MB quantity as described previously.

### 2.7 Spore detection in laid eggs

Eggs were collected from a large cage of MB positive Z line females in a 60 ml pot containing 30 ml distilled water overnight on day three post blood feeding. On day four the pot was removed from the cage three hours prior to processing to allow all viable recently laid eggs to melanize prior to being transferred onto PolysineTM Adhesion Microscope Slides (Epredia – J2800AMNZ) for staining. First, two slides of 40, regularly spaced, white eggs were prepared (directly from the egg pot) by transferring each white egg in 1 µL of the water onto the slide using a micropipette (having cut ∼1 mm off of the end of the tip with a scalpel) and crushing it with an acupuncture needle (length 70 mm, diameter 0.4 mm) to release the egg contents into the water. As the white eggs are fragile, some spores are released into the water during/after laying, so a slide with 40 spots of 1 µL of water from the egg pot was made, intentionally collecting from different areas of the pot each time. Finally, black eggs were carefully collected using a 10/0 paint brush into a 100 µM MACS SmartStrainer (Miltenyi Biotec – #130-098-463) and rinsed well with distilled water to wash off any spores that may have been on the outside of the egg. Two slides of 40 black eggs were made by transferring each egg with a 10/0 paintbrush into 1 µL distilled water (which had already been added onto the slide) and crushed using an acupuncture needle to release the egg contents into the water. The water was allowed to evaporate entirely on each slide before it was incubated at 60 °C for one hour on a hot plate. Each slide was then stained using the “Triple stain” method described by Parry et al. (2025) replacing SYTOX green with 1:1000 NucSpot 470 (Biotium – #40083) for 30 min to stain nuclei.

A single comparable field of view per egg/water drop was imaged with a 10X objective on a LEICA LAS X DM8 microscope with 470, 527 and 700 nM filter cubes and a transmitted light-differential interference contrast (TL-DIC) channel with exposure and gain settings for each channel identical for all images. A manually adjusted z-stack (Z then lambda, 2 µM between slices) was acquired to account for the variation in focal point between the channels.

Maximal intensity projections of all slices for each image were produced in LAS X office software. The number of spore clusters (spore cluster was defined as a 4-10 µM diameter group of around eight objects, with signal in all three fluorescent channels) and presence/absence of spore clusters within the egg husk were counted. When spores were present in the egg husk accurate counting was often restricted; therefore, the number of spore clusters per egg is the minimum. Estimated marginal means with Bonferroni correction was used to compare the number of spore clusters for white eggs, black eggs and water. A representative example for each group was selected for presentation. Each image stack was cropped to reduce the number of frames out of focus and remove the TL-DIC channel, then deconvolved (1 iteration, blind), and maximal intensity projected. One unprocessed image was manually selected from the corresponding TL-DIC z-stack for each group. A panel figure was generated in QuickFigures (Mazo, 2021) of these representative images (which were x-axis cropped) ensuring min/max levels were kept consistent between panels of the same channel. All raw images are deposited on BioImage Archive (Accession: S-BIAD2518).

## 3 Results

### 3.1 MB colony establishment and maintenance

The process of establishing four stable colonies carrying MB is depicted in Fig. 1. We successfully selected one colony named ‘E+’ (Fig. 1) which was derived solely from Village 7 in Valleé du Kou (VK7), other than introduction of laboratory adapted wild type *An. gambiae s*.*l* “G3” males at generation (G) 1. It became stable in terms of high levels of MB transmission at G7. E+ is a mixed line, with more than 50% of individuals producing both *An. gambiae s*.*s* and *An. coluzzii* fragments during a SINE200 PCR (Supplementary Fig. S5) (Santolamazza et al., 2008).

A line from Soumousso village also required addition of laboratory-adapted wild type *An. Gambiae s*.*l* “G3” males at G2. At G6 the colony numbers had declined sufficiently such that it was necessary to combine progeny from two VK7 and one Soumousso strongly MB positive females. The resulting increase in MB prevalence permitted the selection of three VK7/Soumousso hybrid colonies, named ‘G+’, ‘Z+’ and ‘BQ+’ (Fig. 1) which were determined to be *An. coluzzii* through SINE200 PCR (Supplementary Fig. S5) (Santolamazza et al., 2008). The BQ+ line was isolated from the G+ and Z+ lines at G6 and was maintained without selection until G9 at which point the prevalence was 83%. Following the selection of positive parents at G6 (79%) and G7 (97%), the G+ and Z+ lines were separated from one another at G8 from the first (96%) and second (97%) oviposition of the same parents respectively (Fig. 1).

A duplex qPCR assay was designed to permit efficient and accurate quantification of relative MB density (Supplementary Fig. S4). This new assay was evaluated for serial dilutions of CTAB extracted DNA from a pool of ten MB positive adult females, STE extracted DNA from a pool of ten G+ adult females and MB-18S plasmid with minimum quantification thresholds of 1:1000 dilution, 1:125 dilution and 1000 copy number respectively. In all cases the reported duplex assay was found to have acceptable efficiency and R2 values for both probes (efficiency = 90-110%, R2>0.98), indicating accurate quantification within the specified range (Supplementary Fig. S3). The inclusion of locked nucleic acids in the probes was necessary to achieve an optimum annealing temperature and improve specificity.

Using this new assay, the four colonies (BQ+, G+, Z+ and E+) have been maintained for two years with only 1-4 selections required to maintain density and prevalence (Supplementary Fig. S6). Some differences have been observed between the four colonies. The G+ line appears to maintain >80% prevalence most consistently. The BQ+ line required biannual selection due to steadily decreasing prevalence and density. The E+ line was selected only once and maintained for over a year at ∼70% and stable density without further selection. The Z+ line required one selection following a steady decline in prevalence and density in the first year but has since remained stable for 8 months at >80% prevalence and consistent density.

### 3.2 MB density varies between life stages

To better understand MB dynamics across the life cycle of *An. coluzzii*, we investigated our well-established Z+ colony (>1 year stable) in more detail. We evaluated the prevalence and relative density of MB in each distinct life stage and sex of the mosquito host. There were no significant differences in prevalence between life stages according to a two-tailed pairwise-proportion test with Bonferroni correction (Fig. 2A and B) but a one-way ANOVA revealed a significant impact on MB density (Supplementary Table S2) so pairwise comparisons were conducted using two-tailed estimated marginal means tests with Bonferroni correction. The highest mean density was observed in eggs, followed by adult females (Fig. 2C) which was significantly higher than in most other stages (Fig. 2D). The greatest variation in density was observed between eggs (var = 2.6) and the least variation observed in pupal stages (var: male = 0.052, female = 0.049). The lowest mean density was observed in 4th-instar larvae, while there were no significant differences in mean density between larval stages (Fig. 2D).

**Fig. 2:**
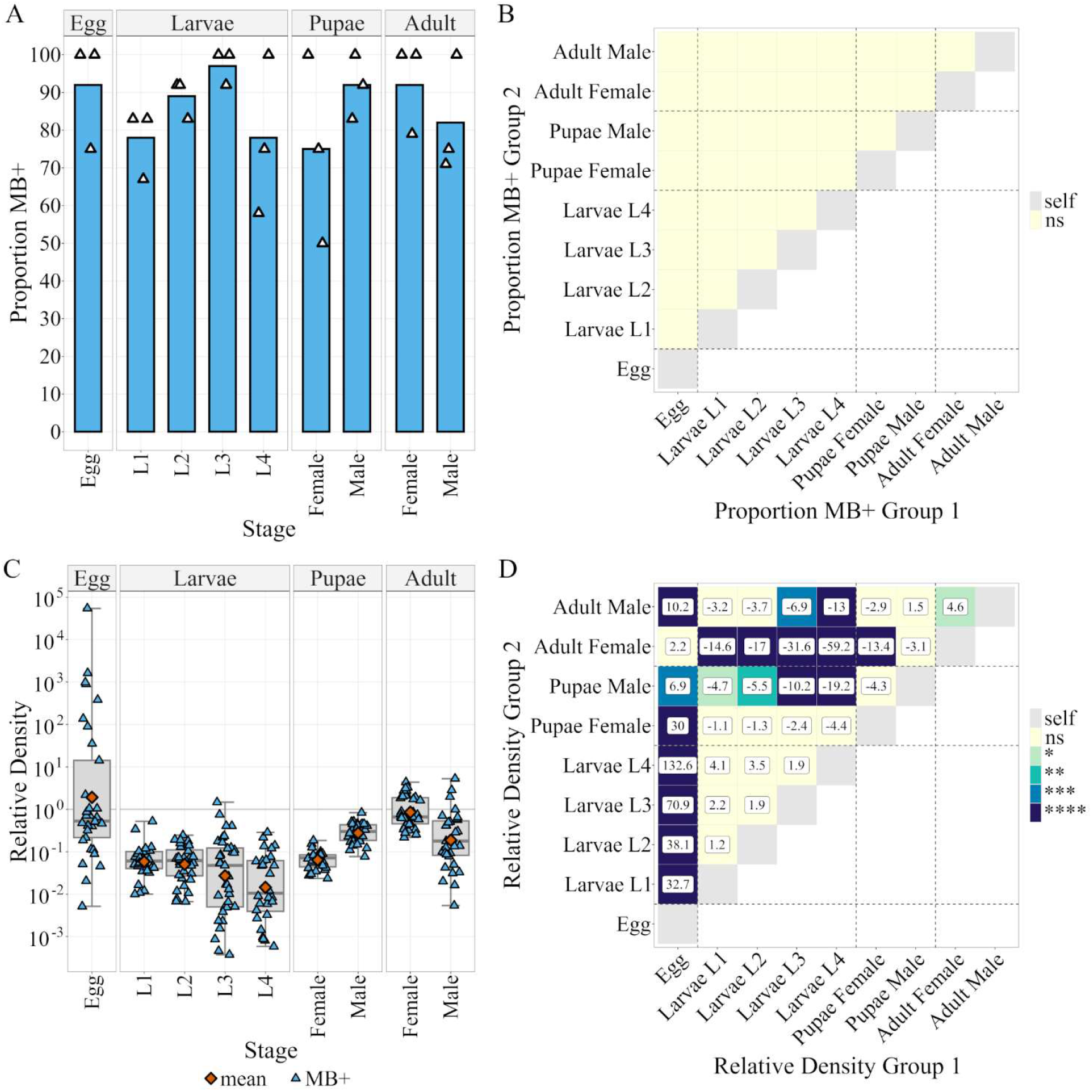
Microsporidia sp.MB prevalence and density in different life stages of Z+ colony. (A) Proportion Microsporidia sp. MB positive mosquitoes at different mosquito life stages as determined by Microsporidia sp. MB TaqMan Cq. n = 36 per life stage (evenly across 3 distinct egg batches). The color indicates p-value: ns > 0.05. (B) Results of a Bonferroni adjusted two-tailed pairwise-proportion test comparing the proportion MB+ in group 1 to group 2. (C) Microsporidia sp. MB density in different life stages relative to *An. gambiae* Elongation Factor Tu, (Relative Density = 2-(CqMB-CqEF)). Boxplots indicate (Min, 25%Q, Median, 75%Q, Max). Orange points indicate the geometric mean otherwise each point reflects an individual mosquito. (D) Results of Bonferroni corrected pairwise estimated marginal means (EMM) test comparing mean log(Relative Density). The values in boxes indicate the fold change in relative density between groups 1 and 2. The ‘-’ symbol indicates that group 2 is larger than group 1. The color indicates p-value: ns > 0.05 > * > 0.01 > ** > 0.001 > *** > 0.0001 > ****.

### 3.3 The prevalence and density of MB are higher in gonadal tissues than in other tissue types

Duplex qPCR was used on several dissected tissues of interest from both female and male adult Z+ mosquitoes. In females there were significant differences in the proportion of MB positive samples between ovaries and other tissues for both virgin and mated unfed females but only between ovaries and salivary glands for mated-blood fed females (Fig. 3A and B). MB prevalence was significantly higher in mated-blood fed carcass (67%) compared to mated-unfed females (11%). When examining mean density, with a two-way ANOVA (mating and blood feeding are aliased coefficients) tissue and the interaction of tissue and ‘mating-blood feeding’ were significant factors (Supplementary Table S2). Pairwise comparisons were conducted using two-tailed estimated marginal means tests with Bonferroni correction. Significantly higher density was observed in ovaries compared to all other tissues (Fig. 3C and D). 4.3-fold higher density (p = 0.003) was observed after blood feeding compared to unfed females when the small number (<7%) of low-density (density < 0.01) ovaries are excluded.

**Fig. 3:**
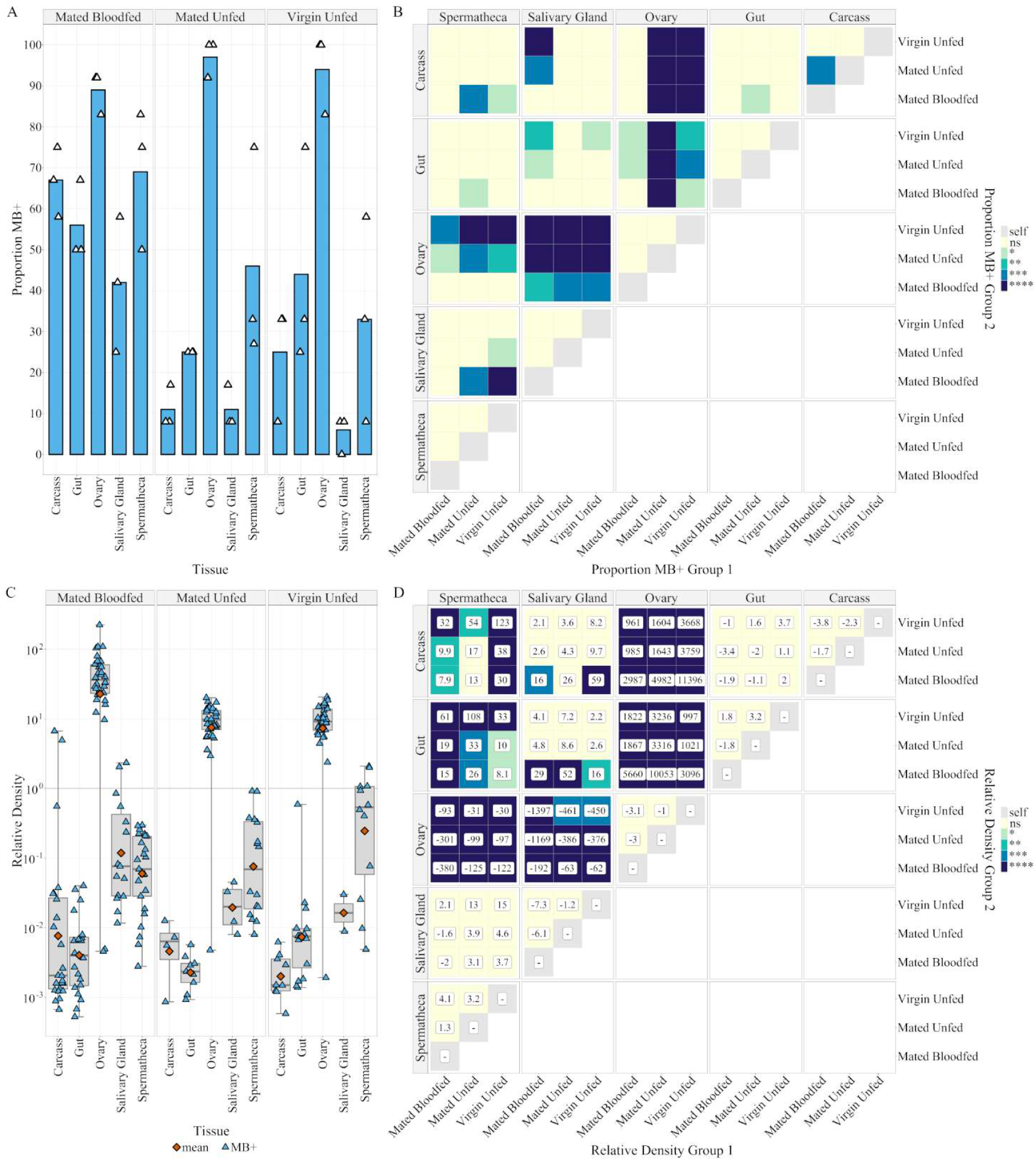
Prevalence and density of Microsporidia sp.MB in female tissues of the Z+ colony. (A) Proportion Microsporidia sp. MB positive samples of different 5-9 day post eclosion adult female tissues as determined by Microsporidia sp. MB TaqMan Cq. n = 36 per tissue (evenly across 3 distinct egg batches). Color indicates p-value: ns > 0.05 > * > 0.01 > ** > 0.001 > *** > 0.0001 > ****. (B) Results of a Bonferroni adjusted two-tailed pairwise-proportion test comparing proportion MB+ in group 1 to group 2. (C) Microsporidia sp. MB Density for 5-9 day post eclosion adult female tissues relative to *An. gambiae* Elongation Factor Tu, (Relative Density = 2-(CqMB-CqEF)). Boxplots indicate (Min, 25%Q, Median, 75%Q, Max). Orange points indicate the geometric mean otherwise each point reflects an individual tissue. (D) Results of Bonferroni corrected pairwise estimated marginal means (EMM) test comparing mean log(Relative Density). Values in boxes indicates the fold change in relative density between groups 1 and 2, ‘-’ symbol indicates that group 2 is larger than group 1. Color indicates p-value: ns > 0.05 > * > 0.01 > ** > 0.001 > *** > 0.0001 > ****.

In males, there was a significant difference in the proportion of MB positives between virgin testes and mated carcass and gut (Fig. 4A and B). When examining density with a two-way ANOVA, tissue type was a significant factor (Supplementary Table S2). Neither mating status nor the interaction between mating and tissue were significant, so pairwise comparisons of tissues were conducted using two-tailed estimated marginal means tests with Bonferroni correction. Mean density in testes was 13.5-fold higher than in gut (p = 8×10^-10^) and 5.7-fold higher than that in the remaining carcass (p = 1.4×10^-5^) (Fig. 4C and D). The density of MB in virgin unfed ovaries was 6.2-fold greater than that in virgin testes.

**Fig. 4:**
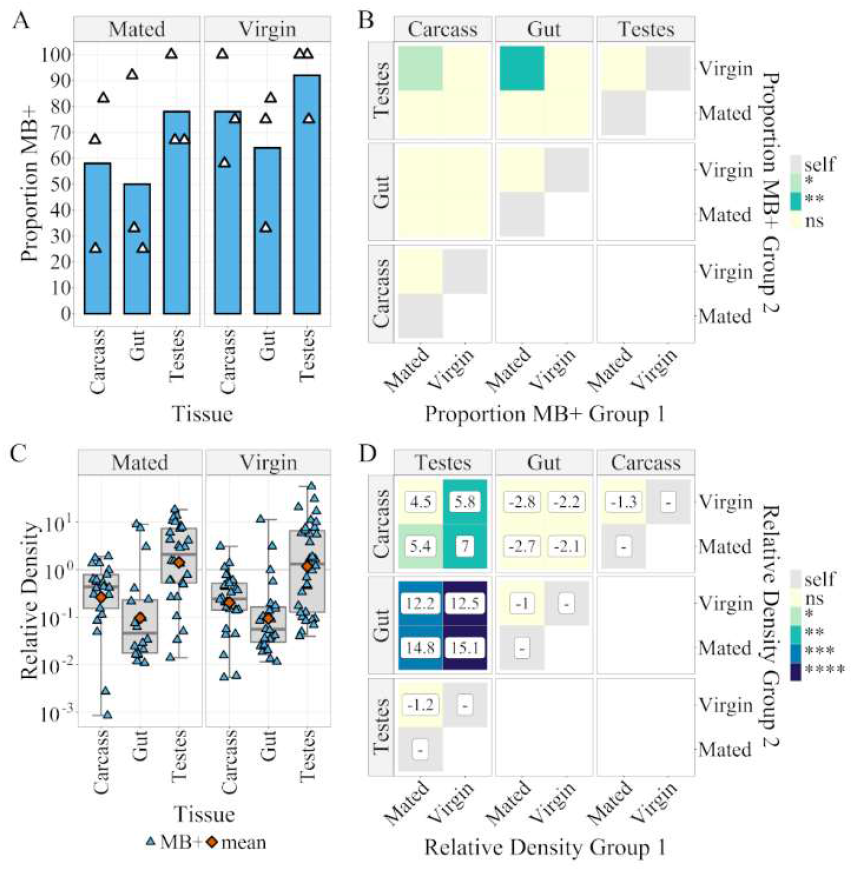
Microsporidia sp.MB prevalence and density in male tissues of the Z+ colony. (A) Proportion Microsporidia sp. MB positive samples of different 5-7 day post eclosion adult male tissues as determined by Microsporidia sp. MB TaqMan Cq. n = 36 per tissue (evenly across 3 distinct egg batches). Color indicates p-value: ns > 0. 05 > * > 0.01 > **. (B) Results of a Bonferroni adjusted two-tailed pairwise-proportion test comparing proportion MB+ in group 1 to group 2. (C) Microsporidia sp. MB Density for 5-7 day post eclosion adult male tissues relative to *An. gambiae* Elongation Factor Tu, (Relative Density = 2-(CqMB-CqEF)). Boxplots indicate (Min, 25%Q, Median, 75%Q, Max). Orange point indicates the geometric mean otherwise each point reflects an individual tissue. (D) The results of Bonferroni corrected pairwise estimated marginal means (EMM) test comparing mean log(Relative Density). The values in boxes indicates the fold change in relative density between groups 1 and 2. The ‘-’ symbol indicates that group 2 is larger than group 1. Color indicates p-value: ns > 0.05 > * > 0.01 > ** > 0.001 > *** > 0.0001 > ****.

### 3.4 Fecundity Assessment

Throughout the rearing process we qualitatively monitored each colony for potential pathogenic effects of MB on its mosquito host including longevity, blood feeding rate and time to pupation. The only identified difference was a possible reduction in the number of eggs laid in positive compared with negative colonies. To investigate this, females of the Z+ and Z-colonies were separated to lay eggs individually. A small percentage (∼7%) of females in the Z+ group were MB negative by duplex qPCR for MB, which, in this experiment, look phenotypically more like the Z-strain than MB positives of the same colony. The laying rates for Z+ (81%) and Z- (80.6%) were very similar (Fig. 5A) with a mixture of normal black eggs and white eggs, which did not undergo normal melanization produced by both MB positives and negatives. A one-way ANOVA, confirmed a significant reduction in median number of eggs per female from 118 for Z-to 104 for Z+ (Fig. 5B, Supplementary Table S2). There was a 13-fold increase in median number of white eggs per female from 1 in Z-to 13 in Z+ which was reflected in an increase in the mean proportion of white eggs per female from 0.8 % in Z-to 12.5 % white eggs in Z+ (Fig. 5B). The median number of black eggs laid by Z+ females (88) is 23% lower than by Z-females (115). Spearman’s correlation test results indicate that increased relative density of MB in positive parent females results in a very weak positive correlation with the number of white eggs (R =0.16, p = 0.045) and that there is no correlation with the number of black eggs (R = -0.07, p = 0.4) or the total (R = -0.05, p = 0.55) number of eggs produced (Supplementary Fig. S7).

**Fig. 5:**
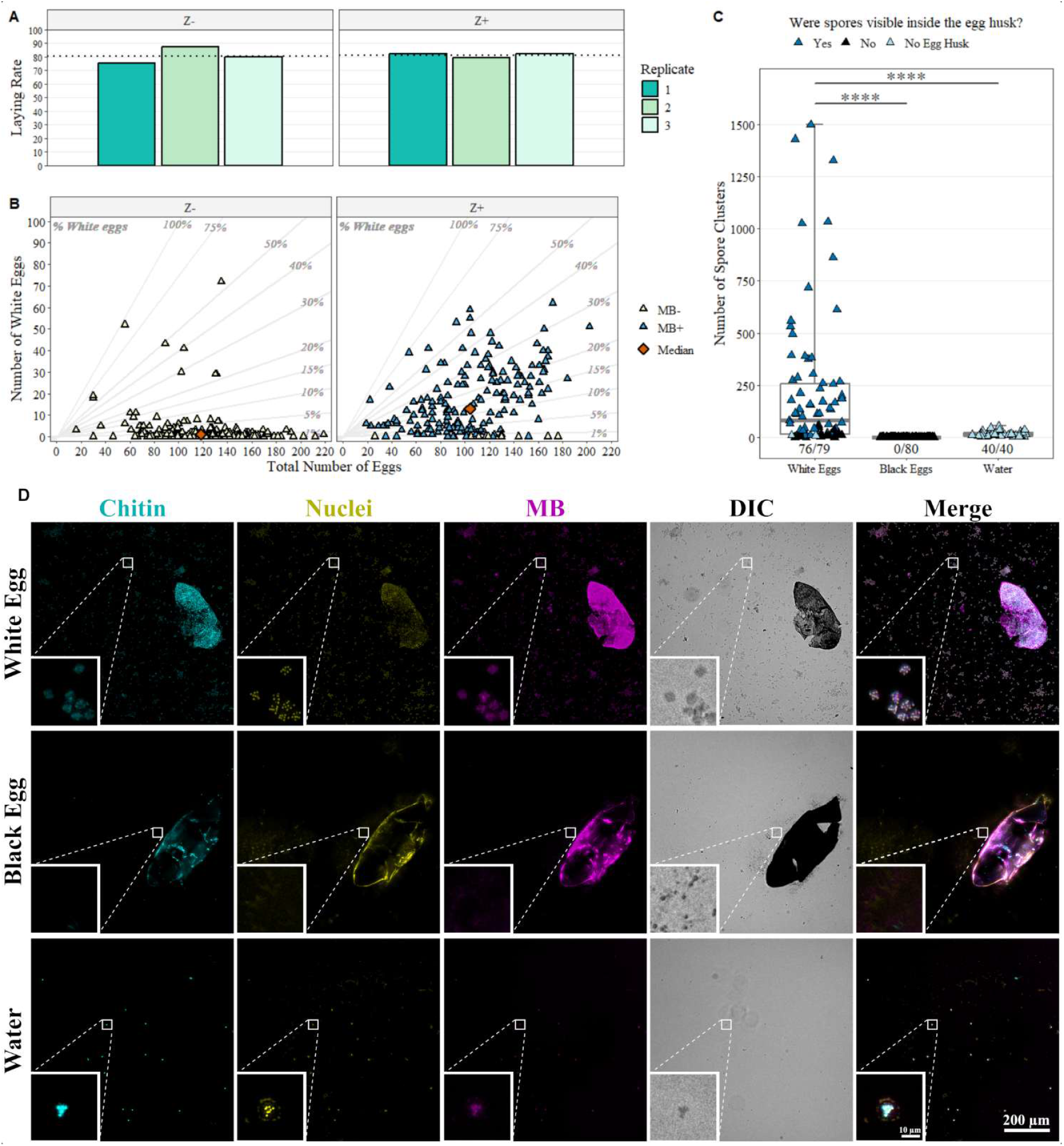
Fecundity is reduced by Microsporidia sp.MB sporogony. (A) Proportion of Z negative (Z-) and positive (Z+) colonies which laid more than 1 egg as part of the fecundity assessment. (B) Number of eggs laid in total against the number of white eggs where each point represents an individual female mosquito. MB+ (blue) and MB- (off-white) were determined by MB duplex qPCR. The median for each group is indicated by an orange diamond and diagonal guidelines indicate the percentage of white eggs. (C) The number of spore clusters found in white and black eggs and the water from the egg pot. “number of eggs with spore clusters / total number of eggs” for each group is indicated at the base of the plot. Asterisks reflect significant estimated marginal means test with Bonferroni correction p-values (****: p<1×10^-5^) for the marked comparison. (D) Representative examples of images collected to evaluate which eggs contained spores (stained with Calcufluor white (chitin - blue), NucSpot 470 (nuclei – yellow) and ATTO 594 MB 18S FISH probes (MB-18S – magenta)). Consistent exposure times and gain settings were used when collecting images. For fluorescent channels, z-stack cropping and post-processing deconvolution of 1 iteration blind were conducted in LAS X office software and maximal intensity projection was conducted in ImageJ. The panel figure was then generated in QuickFigures ensuring that min/max levels for each channel are consistent between panels. Small boxes highlight presence or absence of spore clusters with 8x scaling (bilinear interpolation). Scale bars are indicated for both large and small boxes. Raw images for the entire dataset are deposited on BioImage Archive (Accession: S-BIAD2518). Included in the archive is a file “Images for paper.lif” within which ‘Scene 34 = White Egg’, ‘Scene 16 = Black Egg’ and ‘Scene 25 = Water’ in this figure and Fig5D are presented both raw and post z-stack cropping and deconvolution.

When examining crushed eggs after laying, spore clusters (defined as a 4-10 µM diameter group of around eight objects, with signal in all three fluorescent channels) were counted surrounding 96% of white eggs (median = 80 clusters) and were visible inside 77% of white egg husks. Significantly fewer spore clusters were counted in the water collected from the egg pot (median = 11 clusters) as confirmed by a two-tailed estimated marginal mean test with Bonferroni correction (p = 1.6×10^-6^). Meanwhile, no spore clusters were observed in or surrounding black eggs, which were washed before crushing (Fig. 5C and D, Supplementary Figure S8).

## 4 Discussion

We have, to our knowledge for the first time, successfully established several stable MB carrying *An. gambiae s*.*l* mosquito colonies by employing molecular identification of female *An. gambiae s*.*l* from Burkina Faso carrying the symbiont and intensive selection of their progeny. This is an important step which will greatly facilitate the study of MB and an essential step if it is to provide a new malaria control approach.

Three previous attempts failed before the successful establishment of four stable MB positive colonies. The key differences in the successful attempts compared with failed attempts were: access to larger initial numbers of larvae carrying MB; addition of male laboratory-adapted wild type *An. gambiae s*.*l* “G3” at G1 (VK7) and G2 (Soumousso) which may have improved mating and blood feeding (not quantified); early identification and isolation of pathogenic fungal infections; improved methods for molecular detection of MB; and selection of “strong” positive samples. However, it should be noted that there were many differences and most of these were indiscriminate or uncontrollable, so it was not possible to definitively determine the most important factor. It should be noted that the addition of “G3” mosquitoes, although necessary to establish the colony, will have impacted the genetic diversity and gene expression of these colonies so there are further experiments ongoing to examine this and establish pure MB positive wild type lines. The colonies we have established can be repeatedly crossed with wild type mosquitoes prior to releases to generate lines carrying MB which have genetic backgrounds matching the mosquitoes in the field.

Owing to issues with inconclusive results when using existing methods, a new endpoint PCR assay (Amp-V2) was used briefly (G2-8) while a new qPCR assay was optimized to be highly selective and more efficient (used from G7 onward). The new probe-based duplex qPCR assay for the quantification of MB relative density substantially improved our ability to detect and quantify MB in a variety of different sample types. This assay also permitted replacement of the slow and low throughput CTAB method for STE buffer and boiling for DNA extraction in most instances. This may also have improved our detection and quantification of spore stages as a similar phase separation-based extraction method to CTAB performed poorly for DNA extraction from spore stages with other microsporidia (Subrungruang et al., 2004). Expanding the extraction methods available combined with the duplex nature of the assay makes the MB duplex qPCR far more efficient than endpoint PCR in terms of both time and expense.

We successfully maintained four distinct colonies with minimal intervention for two years, at high MB densities and with standard rearing methods, which has facilitated examination of variation in MB over time. The MB prevalence and density tend to decline more readily in the BQ+ line than in the other lines and thus required biannual selection, whereas in the other lines these tend to remain high with little intervention. This is to our knowledge the first report of successful maintenance of stable mosquito colonies carrying MB at high-density. The ability to maintain MB colonies in this way leads to increased optimism that the prevalence and density of MB can be increased in field populations for malaria control. The possible routes of dissemination of MB are release of live mosquitoes carrying MB in the field, relying initially on vertical transmission, or distribution of MB spores, which would rely on horizonal transmission in early stages. For both approaches, efficient mass rearing of live mosquitoes carrying MB at high densities is likely to be essential.

MB has been shown to block *Plasmodium* falciparum in *An. arabiensis* (Herren et al., 2020) but this has yet to be definitively confirmed in *An. gambiae* carrying MB. However, unpublished preliminary data from experimental challenges indicate greatly decreased susceptibility to *P. falciparum* (from patient blood in Burkina Faso) in MB positive females from the BQ+, Z+ and G+ colonies described here (Sare. S et al, unpublished data). The mechanism underlying the blocking effect has not yet been determined, but investigating the interactions of MB and its mosquito host is crucial to understanding and predicting how MB will spread in field populations. As a first step, we examined the differences in the prevalence and density of MB between different life stages and different tissues. The differences in density observed in different life stages are consistent with those observed through microscopic analysis by (Parry et al., 2025).The higher density MB in adult females compared to L4 larvae and adult males, which we have observed is supported by similar results from qPCR and microscopy in *An. arabiensis* (Makhulu et al., 2024) though in other qPCR results from *An. arabiensis* this difference is less distinct (Herren et al., 2020). These results support the theory that vertical transmission to progeny and horizonal transmission through eggs containing spores are important as these stages harbor the highest densities of MB, so are the most likely stages for onward transmission.

We detected MB in all female tissues, but the prevalence and density were much higher in ovaries compared to other tissues, which is similar to the findings in *An. arabiensis* (Makhulu et al., 2024). However, the presence of MB in the gut, carcass and salivary glands was not observed in microscopic analysis of females from the same colony (Parry et al., 2025). This is likely due to lower throughput and sensitivity of detecting MB using FISH and microscopy of tissue sections. The presence of MB in the female gut and carcass has been observed in *An. arabiensis* using qPCR also at lower prevalence and density than in ovaries and this has been observed in microscopic analysis for a small number of *An. arabiensis* gut samples (Makhulu et al., 2024). Additionally, we observed an increase of ∼4.3-fold in MB density in the ovaries of high-density individuals (> 90% of positive females) following blood feeding which is supported by microscopic analysis of the same strain (Parry et al., 2025). Fluorescence and electron microscopy revealed the development of spores inside some ovarioles after blood feeding, which is likely the cause of the increased relative density. Spores have yet to be confirmed in *An. arabiensis*, although a similar but large increase in MB density following a blood meal has been observed (Makhulu et al., 2024). This difference in quantification could be attributable to our examination at three-days compared with five-days post blood feeding in *An. arabiensis* and it is most likely also the result of sporogony. Interestingly, blood feeding also triggers expansion of tissue tropism with significant increases in MB prevalence in the carcass and salivary glands. This diversification could be an important factor in how MB increases the refractoriness of the host to *Plasmodium* infection, as the change in distribution occurs in parallel with the establishment of the malaria parasite in the gut. Experiments examining the effect of blood feeding in *An. arabiensis* only included comparison of gut and ovary tissues, with a slight reduction in prevalence in gut (Makhulu et al., 2024), where we see an insignificant slight increase in that case.

As in female tissues no effect of mating on MB distribution was observed in male tissues, and similar results were observed to those from *An. arabiensis*, which has the highest prevalence and density in gonadal tissue (Makhulu et al., 2024). In both mosquito species a small number of gut samples were found to have high relative densities of MB, but with higher relative density of MB in the carcass than was observed in *An. arabiensis* (Makhulu et al., 2024) though there have only been limited observations of this using microscopy thus far. These results support potential horizontal (from male to female) and transovarial (transfer from seminal fluid to eggs during oviposition) transmission routes.

Although we were able to maintain high prevalence colonies without difficulty, MB has a negative impact on fecundity in the form of an increased number and proportion of white (non-melanized so presumed unviable) eggs, a reduction in melanized eggs and in total egg number. Fluorescent in situ hybridization (FISH), nuclear and calcofluor staining of sections comparing blood fed to unfed females in the same Z+ colony as was examined here demonstrated that blood feeding coincides with sporogony in a small percentage of developing eggs and confirmed presence of spores at day three post blood feeding using electron microscopy (Parry et al., 2025). From hematoxylin and eosin imaging it seems unlikely that spore-containing eggs contain the required components to melanize at the time of laying. Thus, we theorized and have now confirmed that most (77%) of the non-melanized eggs contained spores and that melanized eggs do not. Due to lack of melanin the white eggs are very fragile and are prone to bursting, which results in the release of spores into the water of the oviposition site ready to infect a new host. With around 50% of white eggs having contained over 100 spore clusters (> 800 spores), this horizonal transmission route is likely an important element of MB’s life cycle.

In any one embryo MB appears to follow one of two developmental routes: multiplication as meront stages in eggs which remain viable leading to vertical transmission, or sporogony in ultimately non-viable eggs leading to potential horizontal transmission. It is possible that already non-viable eggs present optimal conditions for sporogony, since vertical transmission of MB will not occur in that case, whereas the production of spores allows a route of transmission. This reduction in fecundity and spore production has not been observed in *An. arabiensis* (Herren et al., 2020) but it is possible that this is a laboratory versus field difference. Alternatively, this may reflect an MB strain difference, or MB may behave differently in these closely related species (*An. coluzzii* and *An. arabiensis*), as has been reported for other species of Microsporidia. For example, *Enterocytozoon hepatopenai* causes hepatopancreatic microsporidiosis in the Asian penaeid shrimp aquaculture industry but is substantially more pathogenic in *Penaeus vannamei* than in *Penaeus monodon* (Chaijarasphong et al., 2021). Although sporogony does not appear to be detrimental to the female host, the potential reduction in progeny observed here is the first report of MB having an associated fitness cost.

When considering MB as a tool for malaria control this reduction in fecundity could have an impact in several ways. First, it is possible that it is significant enough to impact MB dissemination and increase the effort required to achieve high frequency if the release of MB carrying mosquitoes is the primary method used. However, the horizontal transmission resulting from spores released from eggs may counteract the negative effect of decreased fecundity to increase MB spread. Further experiments examining transmission will be required to distinguish between these possibilities and determine whether any fecundity reduction occurs under field conditions in *An. coluzzii*. In addition, once a high population frequency is achieved, the reduction in fecundity would provide a small reduction in population size / carrying capacity. Finally, we have yet to confirm whether these spores can directly infect *An. gambiae s*.*l* but well understood lifecycles of other mosquito infecting microsporidia would suggest that larval stages consuming the spores released into the oviposition site is a likely transmission route (Becnel, 1994). However, it should be noted that microsporidia showing high levels of horizontal transfer in mosquitoes are more pathogenic than MB. Alternatively (or additionally), it is possible that these spores are part of a digenetic life cycle in which they infect an additional host. The use of spores to disseminate MB is a very attractive possibility as it could be more cost effective and less labor-intensive than the release of live mosquitoes. However, this approach has not yet been tested for the distribution of organisms in the field, whereas release of live adults has proven successful for the dissemination of *Wolbachia* symbionts (Hoffmann et al., 2014, Nazni et al., 2019, O’Neill et al., 2019, Velez et al., 2023, Simmons et al., 2024).

We have successfully established several related *An. gambiae s*.*l* colonies carrying MB at high prevalence and density and have maintained these colonies for two years with minimal intervention. Primary localization of MB is in the gonadal tissue of both males and females, which is consistent with high levels of likely vertical transmission to progeny. The presence of MB in other female tissues could be contributing to *Plasmodium* blocking in ways we do not yet understand particularly given the change in tropism in response to a blood meal. The reduction in fecundity observed is the result of sporogony occurring in some eggs. This may impact control programmes using MB though it is not yet clear what this impact will be or whether it will be positive or negative. Our work is a significant step forward in improving the feasibility of MB as a tool for malaria control and will also considerably facilitate further research on the symbiont to improve understanding of its biology.

## Abbreviations

Cq: Cycle threshold
CTAB: Cetyltrimethylammonium bromide
EF: Elongation factor
FISH: Fluorescent in-situ hybridization
G: Generation
“G3”: Anopheles gambiae G3 strain
L1-L4: First through fourth instar larvae
MB: Microsporidia sp. MB
NCBI: National Centre for Biological Information
SINE200: Short interspersed element-200 bp
STE: Sodium chloride-tris-EDTA
TL-DIC: Transmitted light-differential interference contrast
VK7: Vallée du Kou village 7

## 7 Author Contributions

- BCP – Conceptualization, Data Curation, Formal analysis, Investigation, Methodology, Validation, Visualization, Writing – original draft, Writing – review & editing.
- DKP – Conceptualization, Data Curation, Formal analysis, Investigation, Methodology, Validation, Visualization, Writing – original draft, Writing – review & editing.
- AIA – Conceptualization, Investigation, Writing – review & editing.
- SI – Investigation, Resources, Writing – review & editing.
- ERSP – Investigation, Writing – review & editing.
- RP – Investigation, Writing – review & editing.
- THA – Investigation, Writing – review & editing.
- EB – Investigation, Writing – review & editing.
- AD – Conceptualization, Funding acquisition, Project administration, Resources, Supervision, Writing – review & editing.
- SPS – Conceptualization, Funding acquisition, Project administration, Supervision, Writing – review & editing.

## 9 Acknowledgments

We would like to thank Dr. Gareth Lycett for providing the *An. gambiae s*.*l* “G3” colony; Dr. Jeremy Herren and his team for providing MB Control DNA and ongoing discussions on MB; Dr. Lilian Mbaisi Ang’ang’o for providing MB 18S Scaffold Sequences and the SBOHVM Histology Research Service (University of Glasgow) for their support with optimizing data collection for imaging mosquito eggs.

## 10 Data Availability Statement

The datasets generated and analyzed for this study can be found in the BioImage Archive (Accession: S-BIAD2518) and NCBI (Accession: PX674407).

