## Supplementary Information for "Laboratory *An. gambiae s.l*. mosquito colonies show sustained high transmission of Microsporidia sp. MB and a small fecundity cost"

| Name | 5’ Mod | Sequence | 3’ Mod |
| --- | --- | --- | --- |
| MB18SF - Herren *et al*, 2020 | NONE | CGCCGGCCGTGAAAAATTTA | NONE |
| MB18SR - Herren *et al*, 2020 | NONE | CCTTGGACGTGGGAGCTATC | NONE |
| SP118_MB_Ampv2_F | NONE | ATAGTATACTCGCAAGAGTG | NONE |
| SP119_MB_Ampv2_R | NONE | CTGTTATAGCCTCTTCCTTC | NONE |
| SP121_MB-18S_F | NONE | CTTGAATGAGTGAGATCTTTAGAC | NONE |
| SP122_MB-18S_R | NONE | TCCATCAGAACATCTTCCTC | NONE |
| SP172_An.gam-EF_F | NONE | AGCACAGCAGAAGGAGGAAC | NONE |
| SP173_An.gam-EF_R | NONE | CCACATGGCCGATAGTACCC | NONE |
| *Ag*EF_Probe | 5HEX | CA +C+AT +T+G+C +AGT | IABkFQ |
| MB-18S_Probe | 6-FAM | AT+C+CTTT+TATT+TGC+TAT+TGT | IABkFQ |
| SP77_S7_F | NONE | TCCTGGAGCTGGAGATGAAC | NONE |
| SP78_S7_R | NONE | GACGGGTCTGTACCTTCTGG | NONE |
| SP79_MB_QPCR1_F | NONE | CCACTGAAACAGGAAGGAAGAGG | NONE |
| SP80_MB_QPCR1_R | NONE | CCTTTTATTTGCTATTGTATTGCGCG | NONE |
| SP75_SINE200_F – Santolamazza *et al,* 2008 | NONE | TCGCCTTAGACCTTGCGTTA | NONE |
| SP76_SINE200_R – Santolamazza *et al,* 2008 | NONE | CGCTTCAAGAATTCGAGATAC | NONE |
| SP35_Microspor-MB_Fwd | NONE | CGCCGGCCGTGAAAAATTTA | NONE |
| SP74_Microspor-Ubiq_rev-alt | NONE | GGTTACCTTGTTACGACTT | NONE |

**Supplementary Table S1 – Primers and Probes Information.** *Table details the sequences and modifications used for the primers and probes used in this manuscript. In sequence column a ‘+’ symbol after a base indicates a locked nucleic acid (LNA) base.*

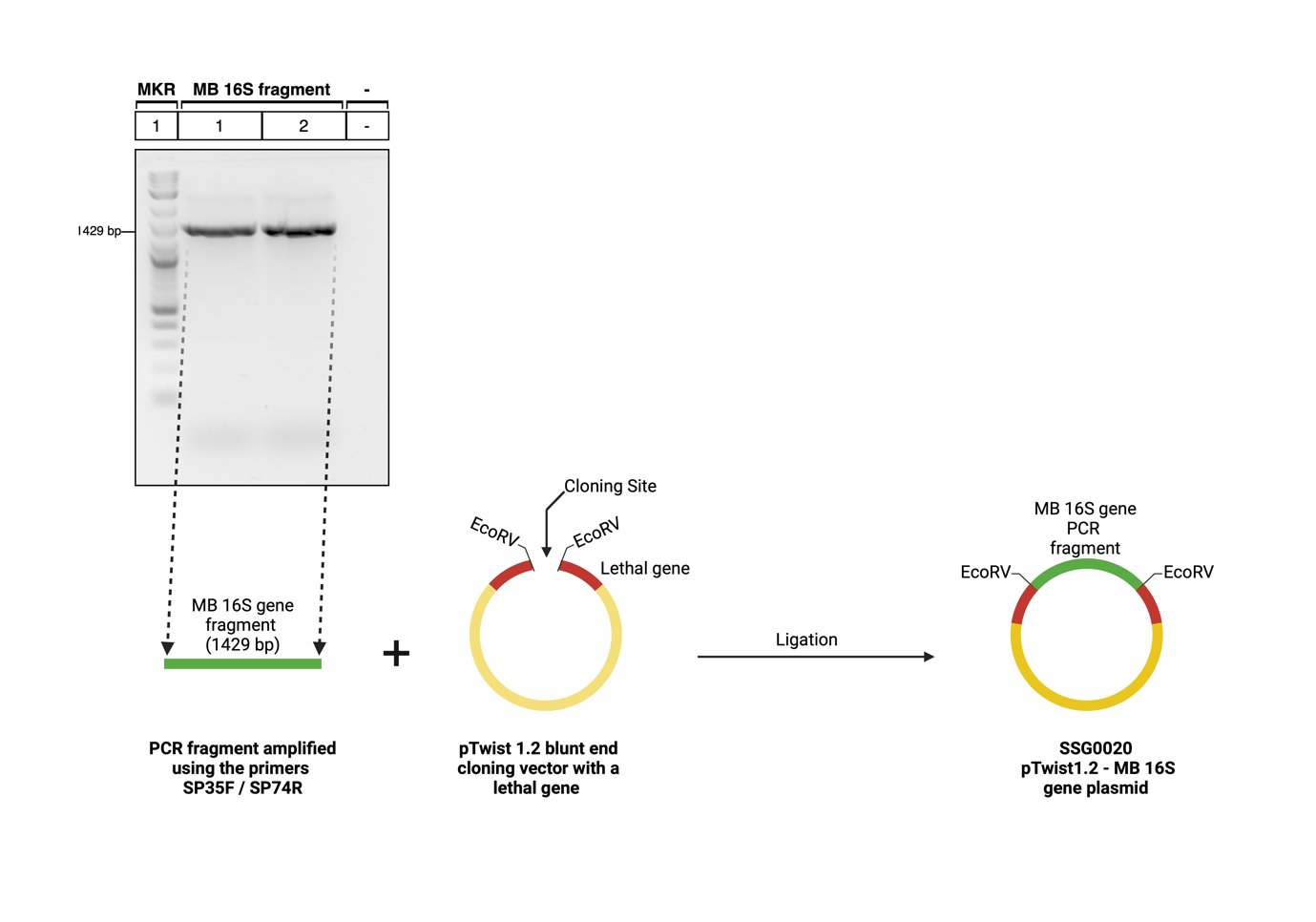

**Supplementary Figure S1 – Cloning of the MB 16S gene fragment into a blunt-end plasmid vector**. *A schematic representation of the cloning strategy used to generate SSG0020 pTwist1.2-MB 16S gene plasmid. Agarose gel analysis of the MB 16S PCR fragment to confirm the fragment length (1429 bp) using 1% agarose gel (top gel image). The purified MB 16S fragment was cloned into the pTwist1.2 blunt-end vector (Thermo Scientific) using T4 DNA ligase (bottom graphics).*

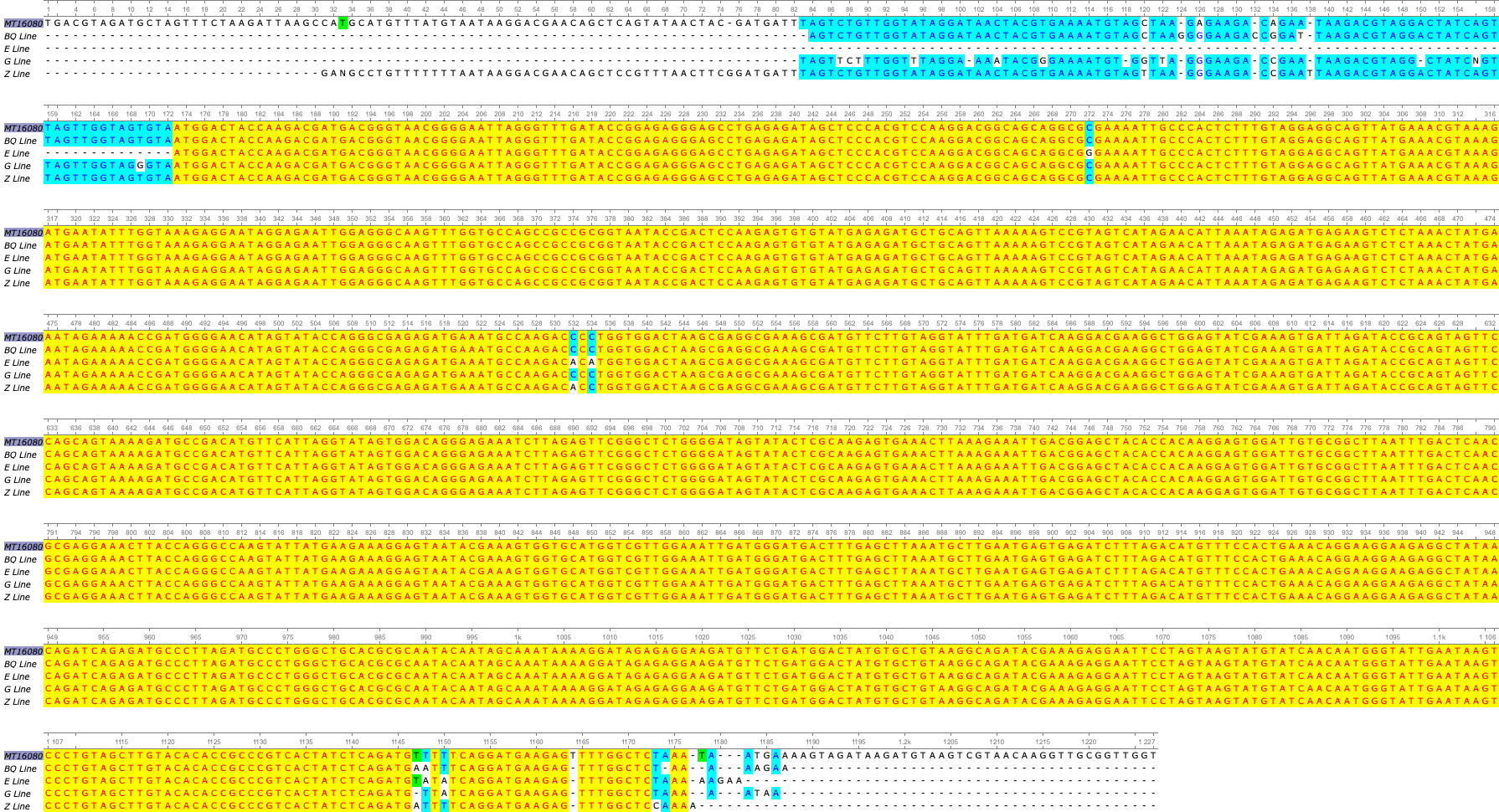

**Supplementary Figure S2 – Multiple sequence alignment of 18S rRNA gene sequences from different MB Lines.** *Only agreements across the different sequences were highlighted and the colours were based on the percentage identity (>50%).*

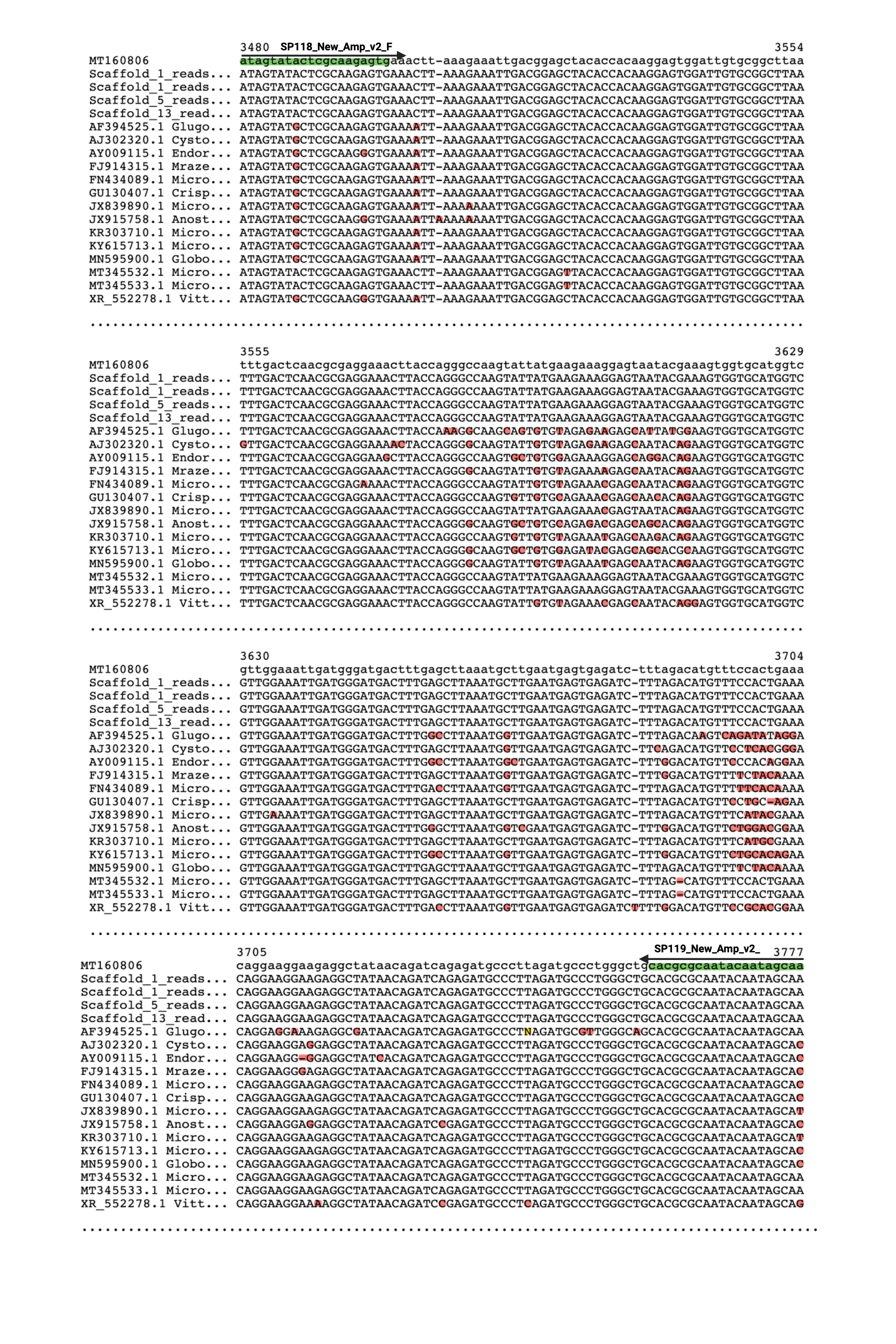

**Supplementary Figure S3 – Development of new 16S end-point assays for Microsporidia sp. MB**

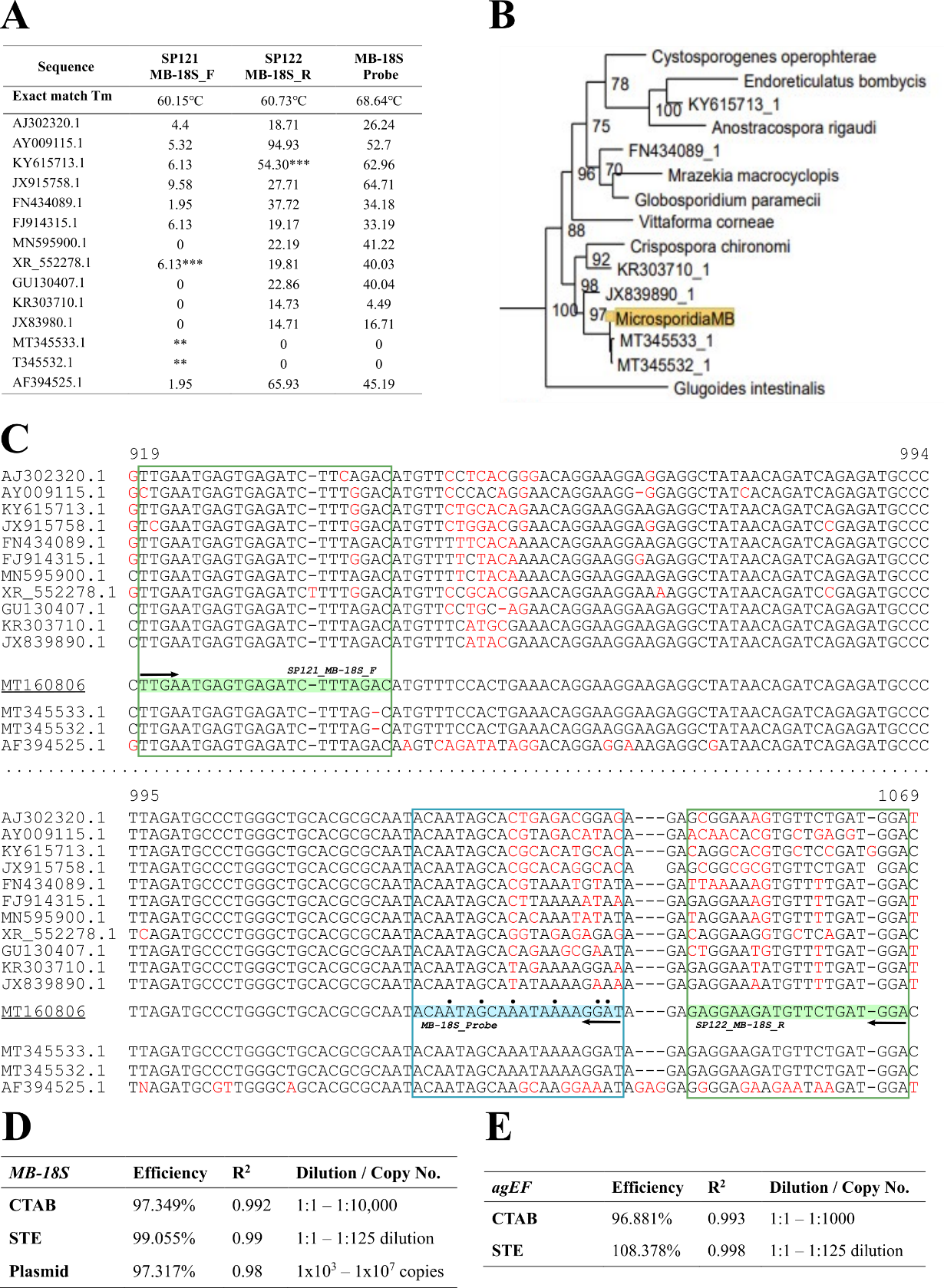

**Supplementary Figure S4** **– TaqMan assay Development and validation.** ***(****A) Indicates the Tm difference between the sequence on the left (neighbouring microsporidian 18S sequences) and the perfect Tm for the primer/probe (>10℃) is considered ideal for selective binding. This is the case for most of the sequences for the reverse primer and probe. (B) A phylogenetic tree of microsporidian 18S sequences, black box indicates the sequences used in the alignment for the design of the TaqMan assay. (C) A fragment of the Microsporidia sp. MB 18S sequence aligned to neighbouring microsporidian 18S sequences indicating differences and the probe and primer locations. Arrows indicate the direction of the sequence and dots above a base indicate that it is a locked nucleic acid in the probe. (D&E) Results from standard curves using MB-18S and AgEF probes respectively*

| **Experiment** | **Model** | **Parameter** | **DFn** | **DFd** | **F** | **p** | **sig** | **ges** |
| --- | --- | --- | --- | --- | --- | --- | --- | --- |
| Life Stage | Log RD ~ Life_Stage | Life Stage | 8 | 273 | 26.534 | 3x10^-30^ | * | 0.437 |
| Female Tissues | Log RD ~ Tissue*MandB | Tissue | 4 | 241 | 214.4 | 3.9x10^-78^ | * | 0.781 |
|  |  | MandB | 2 | 241 | 2.965 | 5.3x10^-2^ |  | 0.024 |
|  |  | Tissue*MandB | 8 | 241 | 2.527 | 1.2x10^-2^ | * | 0.077 |
| Male Tissues | Log RD ~ Tissue*Mated | Tissue | 2 | 145 | 24.488 | 6.9x10^-10^ | * | 0.252 |
|  |  | Mated | 1 | 145 | 0.243 | 0.62 |  | 0.002 |
|  |  | Tissue:Mated | 2 | 145 | 0.041 | 0.96 |  | 5.7x10^-4^ |
| Fecundity | Total Eggs ~ Strain | Strain | 1 | 360 | 9.1 | 0.003 | * | 0.025 |
|  | Black Eggs ~ Strain | Strain | 1 | 360 | 44.591 | 9.2x10^-11^ | * | 0.11 |
|  | White Eggs ~ Strain | Strain | 1 | 360 | 106.44 | 4.9x10^-22^ | * | 0.228 |
| Spore counts | No. Clusters ~ Group | Group | 2 | 196 | 24.338 | 3.6x10^-10^ | * | 0.199 |

**Supplementary Table S2 – ANOVA Outcome Statistics.** *Results were generated using the anova_test() function (*[*https://rpkgs.datanovia.com/rstatix/reference/anova_test.html*](https://rpkgs.datanovia.com/rstatix/reference/anova_test.html)*) of the rstatix package v0.7.2 (reported values are defined in the* *anova_summary() function values -* [*https://search.r-project.org/CRAN/refmans/rstatix/html/anova_summary.html*](https://search.r-project.org/CRAN/refmans/rstatix/html/anova_summary.html)*). RD = relative density MB-18S to AgEF (*2^-ΔCq^*). ‘MandB’ = the combination of Mating and blood feeding status as these parameters are aliased coefficients. Group = the variable denoting white eggs, black eggs or water. * = a significant result where p<0.05.*

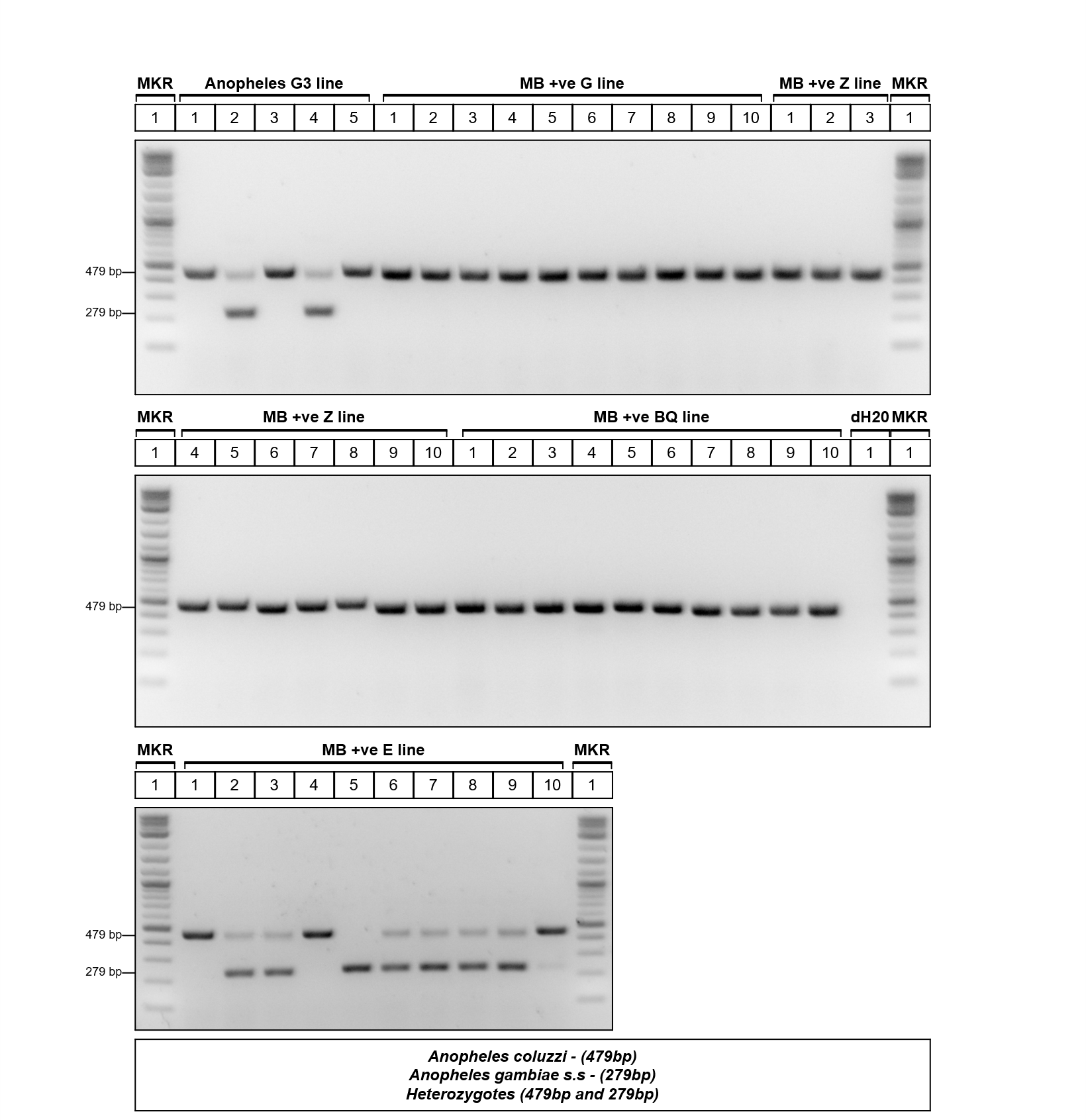

**Supplementary Figure S5 – Agarose gel electrophoresis of SINE200 Diagnostic PCRs in Anopheles G3 line and Microsporidia sp. MB positive lab colonies**. *MKR – NEB Quick-load 1kb DNA ladder. Anopheles G3 – Anopheles gambiae G3 strain (5 samples); MB +ve G, Z, BQ & E line – Microsporidia sp.* MB *positive lab colonies (10 samples each); Anopheles coluzzii produces a single band of 479 bp. Anopheles gambiae s.s produces a single band of 279 bp, and heterozygotes produce both bands.*

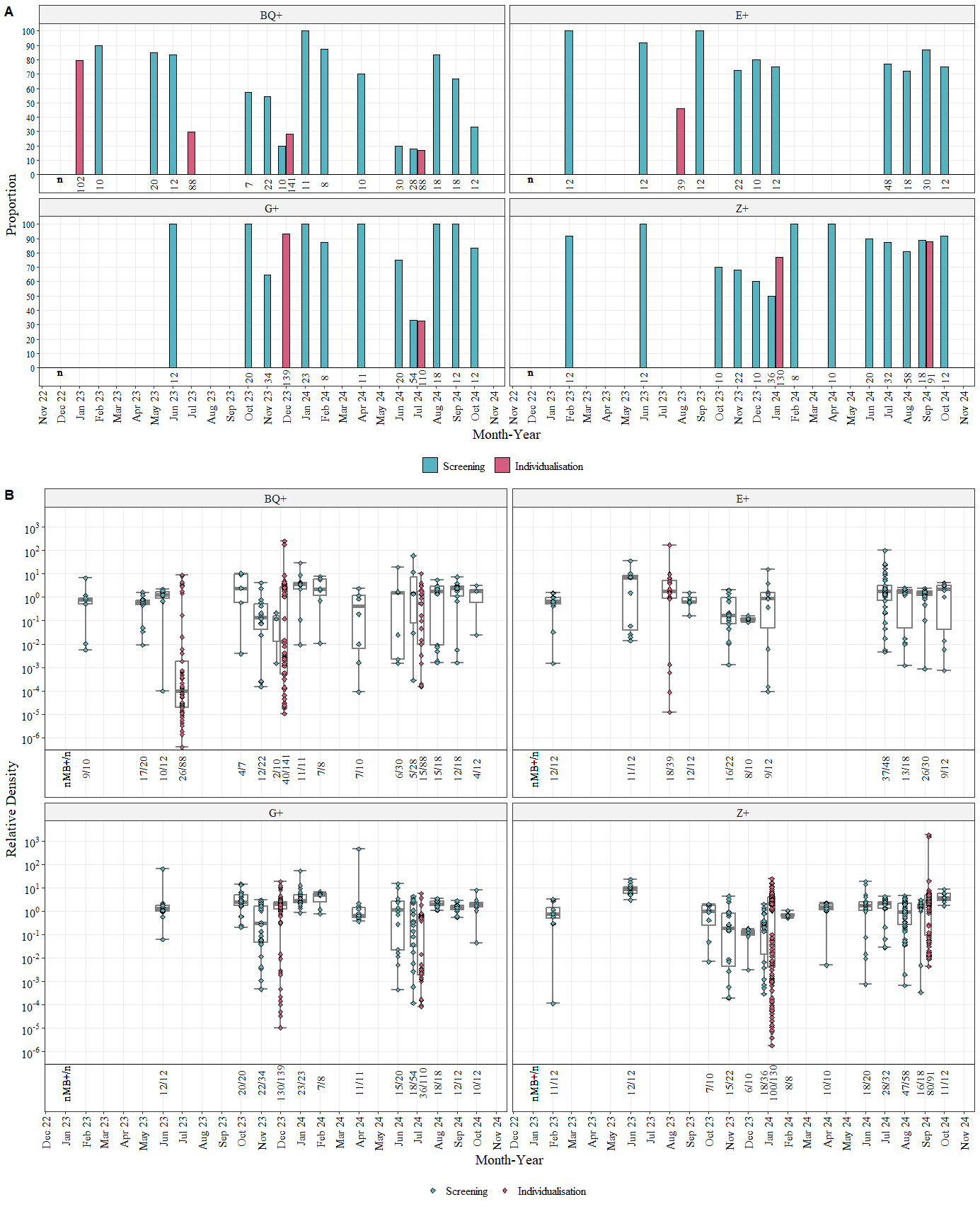

**Supplementary Figure S6 – Ongoing colony maintenance: prevalence and density for the 4 positive colonies we have in our insectary.** *(A) Proportion Microsporidia sp. MB positive samples as determined by Microsporidia sp. MB TaqMan Cq, for colony screens (blue) and individualisations with removal of progeny from Microsporidia sp. MB negative mothers (pink) over 2 years. Facets show the results for the four main colonies we have established in our laboratory. (B) Microsporidia sp. MB Density for different life stages relative to An. gambiae Elongation Factor Tu, (Relative Density = 2^-(CqMB-CqEF)^). Points indicate individual mosquito samples and boxplots indicate (Min, 25%Q, Median, 75%Q, Max).*

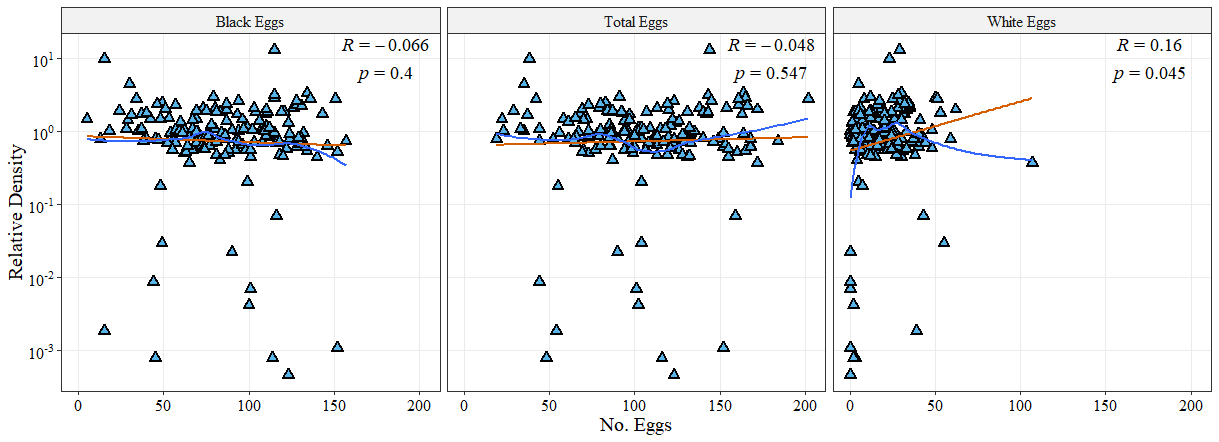

**Supplementary Figure S7 – Egg Type laid by Relative Microsporidia sp. MB Density.** *Microsporidia sp. MB Density in adult females post-oviposition relative to An. gambiae Elongation Factor Tu, (Relative Density = 2^-(CqMB-CqEF)^) compared to the number of black, white and total eggs that individual laid. The orange lines represent the linear model estimate and the blue lines indicate a loess model estimate. Spearmans correlation test R and p values were generated using sm_statCorr().*
