## Supplementary Information 2 for "Laboratory *An. gambiae s.l*. mosquito colonies show sustained high transmission of Microsporidia sp. MB and a small fecundity cost"

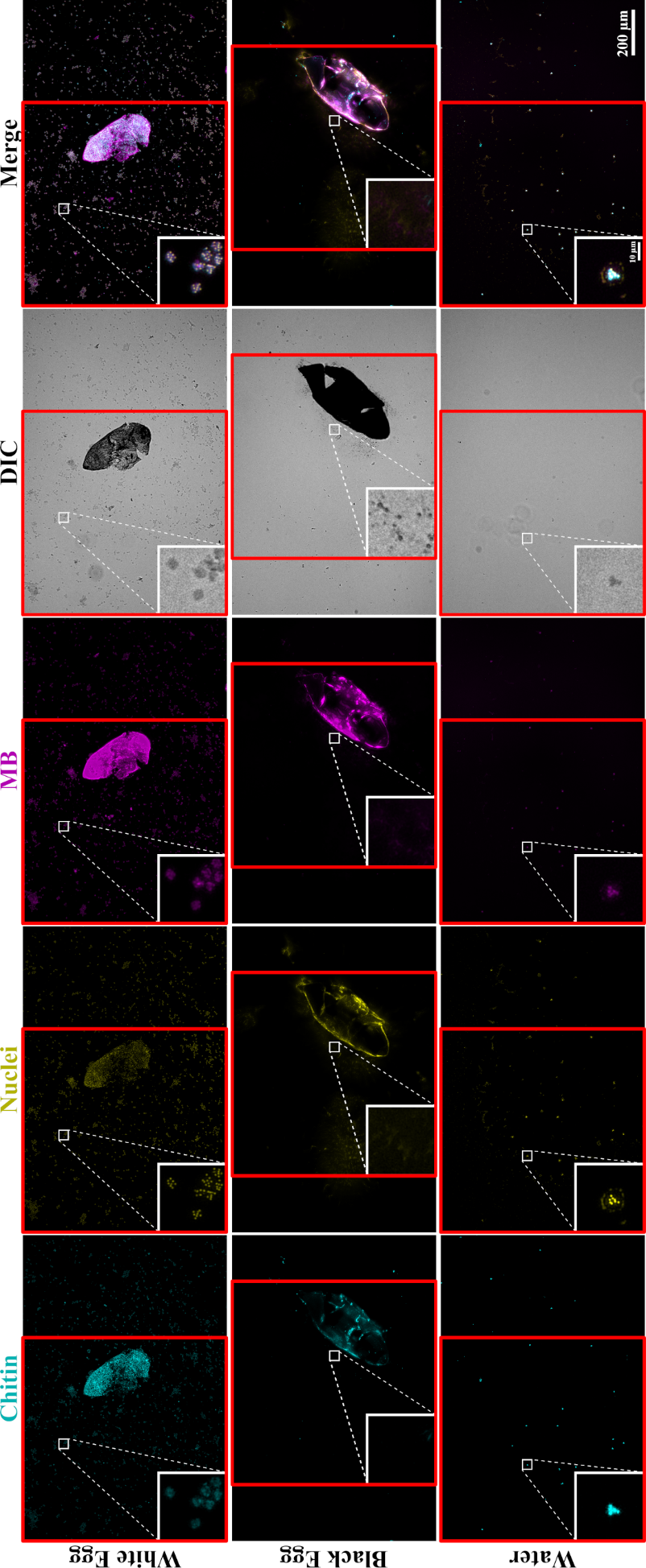

**Supplementary Figure S8 – Uncropped panels for Fig5D indicating cropping regions.** *Representative examples of images collected to evaluate which eggs contained spores (stained with Calcufluor white (chitin - blue), NucSpot 470 (nuclei – yellow) and ATTO 594 MB 18S FISH probes (MB-18S – magenta)). Consistent exposure times and gain settings were used when collecting images. For fluorescent channels, cropping of z-stacks and post-processing deconvolution of 1 iteration blind were conducted in LAS X office software and maximal intensity projection was conducted in ImageJ. The panel figure was then generated in QuickFigures ensuring that min/max levels for each channel are consistent between panels. Small boxes highlight presence or absence of spore clusters with 8x scaling (bilinear interpolation). Scale bars are indicated for both large and small boxes. Red boxes indicate where the images were cropped for ease of presentation in Fig5D. Raw images for the entire dataset are deposited on BioImage Archive (Accession: S-BIAD2518). Included in the archive is a file “Images for paper.lif” within which ‘Scene 34 = White Egg’, ‘Scene 16 = Black Egg’ and ‘Scene 25 = Water’ in this figure and Fig5D are presented both raw and post z-stack cropping and deconvolution.*
